# Tissue resident memory CD8+ T cells are present but not critical for demyelination and neurodegeneration in a mouse model of multiple system atrophy

**DOI:** 10.1101/2024.06.02.597035

**Authors:** Nicole J. Corbin-Stein, Gabrielle M. Childers, Jhodi M. Webster, Asta Zane, Ya-Ting Yang, Md Akkas Ali, Ivette M. Sandoval, Fredric P. Manfredsson, Jeffrey H. Kordower, Daniel J. Tyrrell, Ashley S. Harms

## Abstract

Multiple system atrophy (MSA) is rare, fast progressing, and fatal synucleinopathy with alpha-synuclein (α-syn) inclusions located within oligodendroglia called glial cytoplasmic inclusions (GCI). Along with GCI pathology there is severe demyelination, neurodegeneration, and neuroinflammation. In post-mortem tissue, there is significant infiltration of CD8+ T cells into the brain parenchyma, however their role in disease progression is unknown. To determine the role of CD8+ T cells, a modified AAV, Olig001-SYN, was used to selectively overexpress α-syn in oligodendrocytes modeling MSA in mice. Four weeks post transduction, we observed significant CD8+ T cell infiltration into the striatum of Olig001-SYN transduced mice recapitulating the CD8+ T cell infiltration observed in post-mortem tissue. To understand the role of CD8+ T cells, a CD8 knockout mice were transduced with Olig001-SYN. Six months post transduction into a mouse lacking CD8+ T cells, demyelination and neurodegeneration were unchanged. Four weeks post transduction, neuroinflammation and demyelination were enhanced in CD8 knockout mice compared to wild type controls. Applying unbiased spectral flow cytometry, CD103+, CD69+, CD44+, CXCR6+, CD8+ T cells were identified when α-syn was present in oligodendrocytes, suggesting the presence of tissue resident memory CD8+ T (Trm) cells during MSA disease progression. This study indicates that CD8+ T cells are not critical in driving MSA pathology but are needed to modulate the neuroinflammation and demyelination response.

## Introduction

Multiple system atrophy (MSA) is a progressive, demyelinating neurodegenerative disease characterized by progressive autonomic decline, parkinsonism and ataxia[8, 25, 34]. MSA is classified as a synucleinopathy due to the presence of alpha synuclein (α-syn) in insoluble glia cytoplasmic inclusions (GCIs) located predominantly in oligodendrocytes[8, 25, 28, 34, 37]. Depending on the areas of degeneration within the brain, MSA can be classified into two subtypes. MSA-Parkinsonian (MSA-P) is the most prevalent occurring in approximately 80% of individual with MSA, with neurodegeneration occurring in the nigrostriatal pathway and within the striatum proper [8, 25]. MSA-cerebellar ataxia (MSA-C) occurs in the remaining 20% of MSA patients, characterized by neurodegeneration in the cerebellum[8, 25]. Although MSA subtypes differ based on areas of neurodegeneration, both subtypes have the α-syn GCI pathology, demyelination, and eventually neurodegeneration. With demyelination, there is also evidence of a robust neuroinflammation response during MSA disease progression[8, 22, 25, 30].

Neuroinflammation is a pathological hallmark in MSA[8, 24, 25, 31]. In PET imaging studies, responding microglia can be detected in the striatum and throughout the basal ganglia of MSA patients during disease progression[13]. Additionally, in MSA post-mortem tissue, microgliosis and astrogliosis are seen in areas of demyelination and neurodegeneration[8, 22, 25]. Furthermore, human leukocyte antigen-DR (HLA-DR), also known as major histocompatibility complex II (MHCII), is present on antigen presenting cells (APCs) within the putamen and substantia nigra[22, 40]. This myeloid activation is also accompanied by the infiltration of peripheral CD4+ and CD8+ T cells[40]. To model MSA, a modified adeno-associated virus (AAV) has been engineered via capsid shuffling to overexpress human a-syn in oligodendrocytes (Olig001-SYN)[3, 27]. In rodents and monkeys, the Olig001-SYN vector faithfully replicate MSA pathology including neuroinflammation, demyelination, and neurodegeneration[26]. This α-syn overexpression in oligodendrocytes results in MHCII expression on APCs and an infiltration of CD4+ and CD8+ T cells, that mimics what is observed in MSA post-mortem tissue[40]. Our group has previously shown that CD4+ T cells producing the proinflammatory cytokine IFNγ are critical in driving disease progression in the Olig001-SYN mouse model[7]. Little is known thought about the function of CD8+ T cells during MSA pathogenesis.

CD8+ T cells, also known as cytotoxic T cells, become activated after encountering a non-self-antigen via major histocompatibility complex I (MHCI)[14, 15]. In response, they produce cytokines (IFNγ and perforin) and granzymes (granzyme b and a) that can induce cytotoxic killing of infected cells[14, 15]. There is growing evidence that CD8+ T cells are present within the central nervous system (CNS) in neurodegenerative diseases such as Multiple Sclerosis (MS), Alzheimer’s disease (AD) and Parkinson Disease (PD)[12, 23, 32, 39]. In MS, CD8+ T cells are present within the white matter lesions and are clonally expanded into tissue resident memory T (Trm) cells[9]. Similarly, CD8+ Trm cells have also been found within the CSF, blood, and brain parenchyma of AD and PD patients, respectively[12, 39]. The role CD8+ T cells in neurodegenerative disease is a new area of investigation, but little is known about the role of CD8+ T cells in MSA.

In this study, we sought to determine the role of CD8+ T cells during MSA disease progression. Using the Olig001-SYN mouse model of MSA, we determined that α-syn overexpression in oligodendrocytes resulted in significant infiltration of CD8+ T cells in the brain parenchyma. The genetic knockout of CD8+ T cells did not change demyelination and neurodegeneration in the Olig001-SYN model at six months post transduction. However, at an intermediate four-week timepoint, enhanced neuroinflammation and demyelination was detected in areas of GCI pathology. Utilizing spectral flow cytometry, we show that the CD8+ T cells in the striatum have an early tissue resident memory phenotype (CD103+, CD69+, CD44+, CXCR6+). These findings identified CD8+ Trm cells during MSA and highlight a possible role of CD8+ T cells in the development of neuroinflammation and demyelination in the Olig001 mouse model of MSA. Understanding the complexity of CD8+ T cells in MSA could identify potential targets for disease modifying treatments.

## Materials and methods

### Mice

Male and female C57BL/6 (#000664 Jackson Laboratories) were used for all experiments and maintained on a congenic background. Additionally, both male and female CD8 knockout mice (#002665 Jackson Laboratories) with a C57BL/6 background were used. All research conducted on animals were approved by the Institutional Animal Care and Use Committee at the University of Alabama at Birmingham (UAB).

### Olig001 vector

The Olig001 vector was created via directed evolution and has been previously characterized by our lab and others[3, 7, 27, 40]. The Olig001 virus is a modified AAV capsid with a >95% tropism for oligodendroglia. The expression of either human α-syn or GFP is controlled by a CBh promoter and bovine growth hormone polyA. Vector was packaged via triple-transfection of 293T cells with plasmids encoding vector genome and helper function, and purified using an iodixanol gradient followed by buffer exchange as previously described[33]. Titers were determined using digital droplet PCR and normalized using modified PBS[4].

### Stereotaxic surgery

Male and female mice aged to 8-12 weeks were anesthetized with isoflurane via a vaporizing instrument provided by the Animal Resource Program at UAB. Mice were injected either unilaterally (Black-Gold, DAB, and immunofluorescences) or bilaterally (flow cytometry) with 2ul of either Olig001-GFP or Olig001-SYN (both 1 ×10^13^ vector genomes (vg)/ml). To mimic MSA-P, Olig001 was injected into the dorsolateral striatum (AP + 0.7 mm, ML +/− 2.0 mm, DV – 2.9 mm.) using a Hamilton syringe on an automatic injecting system at a rate of 0.5 μl/min. After the injection was complete, the needle was left in for an additional 2 minutes. The needle was slowly removed over the course of 2 minutes. All surgical protocols and aftercare were followed and approved by the Institutional Animal Care and Use Committee (IACUC) at the University of Alabama at Birmingham.

### Immunohistochemistry tissue preparation and staining

At four weeks and six months post Olig001 transduction, mice were anesthetized and underwent transcardial perfusion with 0.01M Phosphate-buffered saline (PBS) pH 7.4, followed by a fixation with 4% paraformaldehyde (in PBS, pH 7.4; PFA). The brain was isolated and incubated in 4% PFA solution at 4°C solution for 4 hours. After fixation, the brains were cryoprotected with a 30% sucrose (in PBS) solution until brains were fully saturated. Brains were frozen and coronally cryosectioned at 40μm on a microtome. Tissue was stored in a 50% glycerol/PBS solution at –20°C.

Free-floating sections were washed in 0.01M tris-buffered solution (TBS; pH 7.4) 3 X for 5 minutes, then a sodium citrate antigen retrieval (0.1M; pH 7.3) process for 30 minutes at 37°C. Sections were blocked in 5% normal serum for 1 hour. Tissue was incubated in 1% serum in TBS-Triton (TBST) primary antibody solution consisting of one of the following antibodies: anti-Iba1 (1:500, WACO), anti-CD3 (1:500, clone 17A2; Thermo Fisher), anti-MHCII (1:500, M5/114.15.2; Thermo Fisher), anti-CD8 (1:500, clone 4SM15; eBioscience), anti-CD4 (1:500, clone RM4-5; Thermo Fisher), overnight at 4°C. Next day, the tissue was washed and put into a 1% serum TBST secondary solution for 2 hours. Sections were mounted onto plus coated glass slides (Fisher Scientific), and cover slipped using hard set mounting medium (Vector Laboratories). Fluorescence images were collected on a Ti2 Nikon microscope using a Ci2 confocal system.

### DAB labeling and quantification

At room temperature, striatal sections were quenched in a 3% hydrogen peroxide/50% methanol in 0.01M TBS (pH 7.4) solution for 5 minutes. Tissues were then washed and incubated in antigen retrieval sodium citrate (pH 6.0) for 30 minutes at 37°C. Striatal sections were blocked in TBST with 5% serum, then incubated with either a pSer129 (1:5000, clone EPI53644; Abcam) or NeuN (clone EPR12763; Abcam) antibody in TBST with 1% serum overnight at 4°C. The following day, sections were incubated with a biotinylated goat anti-rat IgG secondary antibody (1:1000, Vector Labs) or anti-rabbit IgG secondary antibody (1:1000, Vector Labs) in a 1% serum TBST solution. According to the manufacturer’s protocol, the R.T.U Vectastain ABC Reagent kit (Vecter Labs) and DAB kits (Vector Labs) were used to develop and visualize the stain. Striatal tissue was mounted onto plus coated slides (Fisher Scientific). Slides were dehydrated in ascending gradient of alcohol solutions and CitriSolv (Decon Laboratories Inc.) and cover slipped with Permount (Electron Microscopy Sciences). Images were taken at 10X on a Zeiss imager M2 brightfield microscope (MBF Biosciences). In Supplemental Figure 1, data was analyzed in ImageJ and the fold change of mean grey value between the ipsi– and contra-lateral side were calculated.

### Black-gold staining

To assessed myelin integrity, 40 μm thick sections were heat fixed onto gelatin-coated slides (Southern Biotech; SLD01-BX) and then stained with the Black-Gold II RTD Myelin Stain Kit (Biosensis) according to the manufacturer’s instructions. Briefly, after heat fixation onto gelatin-coated slides, tissue was washed with DI water in glass coplin jars for 2 minutes. Slides were then incubated in the Black-Gold solution at 65°C in the dark for approximately 12 minutes. Slides were rinsed with DI water for 2 minutes and fixed with sodium thiosulfate. Afterwards, slides were rinsed with DI water and dehydrated/cover slipped with the methods described in the DAB staining section.

### Mononuclear cell sorting and flow cytometry

Four weeks post Olig001 transduction, mice were anesthetized and perfused with 0.01M PBS pH 7.4. Brain and other tissues were removed and the striatum was dissected. Brain tissues were triturated and digested with 1 mg/mL Collagenase IV (Sigma) diluted in RPMI 1640 with 10% heat inactivated fetal bovine serum, 1% glutamine (Sigma), and 1% Penicillin– Streptomycin (Sigma). Afterwards, samples were passed through a 70 uM filter. Mononuclear cells were then separated with a 30/70% percoll gradient (GE).

Cells were isolated and blocked with anti-Fcy receptor (1:100; BD Biosciences) before being labeled with the following fluorescent-conjugated antibodies: CD45 (clone 30-F11; eBioscience), CD11b (clone M1/70; BioLegend), MHCII (clone M5/114.15.2; BioLegend), MHCI (clone 28-8-6; BioLegend), Ly6C (clone HK1.4; BioLegend), CD4 (clone GK1.5; BioLegend), CD8a (clone 53.6.7; BioLegend), CD27 (clone g.3A10; BioLegend), CD69 (clone H1.2F3; BioLegend), TCRbeta (clone 37.51; BioLegend), PD-1 (clone 29F.1A12; BioLegend), KLRG1 (clone 2F1; BioLegend), CD62L (clone MEL-14; BioLegend), CXCR3 (clone 173; BioLegend), CCR5 (clone 7A4, BioLegend), CD103 (clone 2E7, BioLegend), CXCR6 (clone SA051D1, BioLegend), CD44 (clone 1M7; BioLegend), anti-PDGFRa (clone APA5; BioLegend). For conventional flow cytometry, a Fixable Near-IR LIVE/DEAD Stain Kit (Invitrogen), fixable viability dye was used to distinguish live cells per manufacturer’s instructions. For experiments using spectral flow cytometry methods, the Fixable Aqua Dead Cell Stain Kit, for 405 nm excitation (Invitrogen) was used to label dead cells.

For intracellular transcription factor labeling of oligodendrocytes, the Foxp3/Transcription Factor Staining Kit (eBioscience) was used accordingly with conjugated antibodies against Olig2 (clone 211F1.1; Sigma Millipore). An Attune Nxt (Thermo Fisher Scientific) or a BD Symphony spectral flow cytometer (BD Sciences) were used to run samples and FlowJo (Tree Star) software were used for analysis. Mean cell count numbers, percentages, and mean fluorescent intensity (MFI) were analyzed with FlowJo software to assess for neuroinflammation.

### Stereological Analysis of Striatum

Six months post Olig001-SYN unilateral injection, tissues were harvested and sectioned as described above. Tissue was stained with NeuN (clone EPR12763; Abcam) and developed with the DAB protocol also described above. The number of NeuN+ cells were estimated on a Zeiss Axio Imager M2 microscope using StereoInvestigator Software (version 2021.1.3; MFB Biosciences). Due to the localization of Olig-001-α-syn injections, the contours used for counting were drawn around the dorsolateral striatum (Figure 5A). To encompass areas of pathology, six serial sections were counted per mouse. Grid size was 300 μm x 300 μm and the counting frame was set at 40×40. After dehydration, the thickness of the sections was determined to be 25 μm. Estimated populations were generated, and a two-way ANOVA and a Tukey post hoc test was used to determine the significance of neuronal counts between groups.

## Statistical Analysis

All graphs and corresponding statistical tests were generated or performed using Prism software (GraphPad) and Jupyter notebook (Python 3). For flow cytometry graphs in Figure 2 and 3, all data points were compared across genotype using a ranked-summed, non-parametric Wilcoxon test (with 95% confidence and p<0.05). Due to the larger sampling groups, flow cytometry graphs in Figure 1 were analyzed with an unpaired student’s t-test with a 95% confidence and p,0.05. A two-way ANOVA was used and Tukey’s Honestly-Significant-Difference (with 95% confidence and p<0.05) was used to determine significance for stereology analysis of WT and CD8 knockout transduced with either Olig001-GFP or SYN.

**Figure 1:**
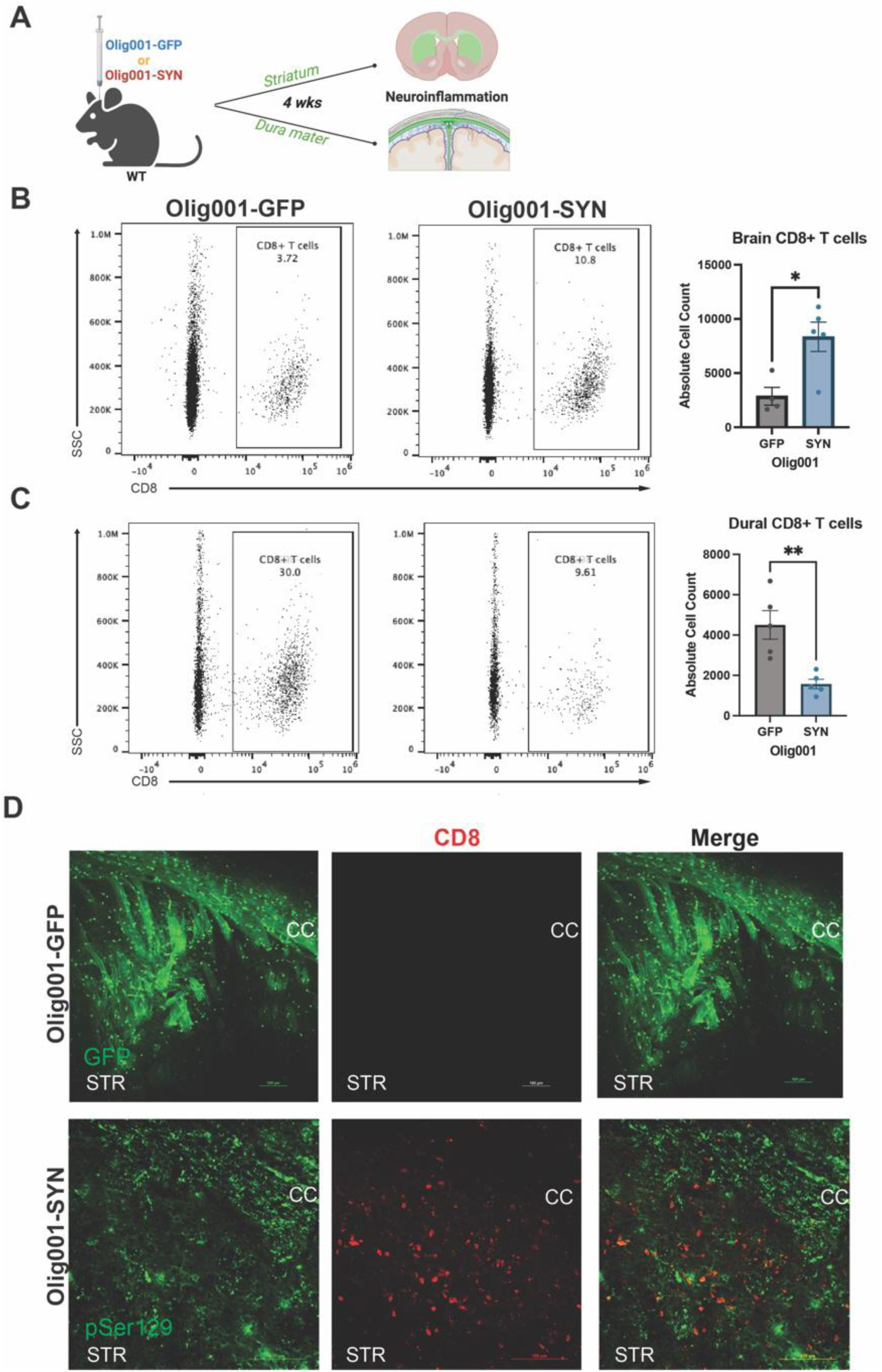
CD8+ T cells are present in the brain and meninges of Olig001-SYN transduced mice. (**A**) 8-12 week old WT mice were bilaterally injected for flow cytometry or unilaterally injected for immunohistochemistry with either Olig001-GFP or Olig001-SYN. After four weeks the striatum, dura mater of the meninges, and leptomeninges were collected and processed for flow cytometry (n=4 or 5; 2 mice were pooled for each sample). (**B**) Representative dot plots from flow cytometry on CD8+ T cells within the striatum of Olig001-GFP and SYN, quantification is to the right of the dot plots. Mean values +/− SEM are plotted, non-parametric Wilcoxon test, *p < 0.05. (**C**) Flow cytometry dot plots of CD8+ T cells from the dura mater of the meninges of Olig001-GFP or SYN transduced mice; quantification is to the right of dot plots. Mean values +/− SEM are plotted, non-parametric Wilcoxon test, **p < 0.01. (**D**) Representative images at 20x magnification of WT mice injected with Olig001-GFP or Olig001-SYN. The corpus callosum is labeled as CC and the striatum is labeled as STR. CD8+ T cells (red) in the parenchyma of striatum within areas were pSer129 (green) in Olig001-SYN injected mice. n=3 for immunohistochemistry staining; scale bar is set at 100 μM. Both male and female mice were used for all experiments.

**Figure 2:**
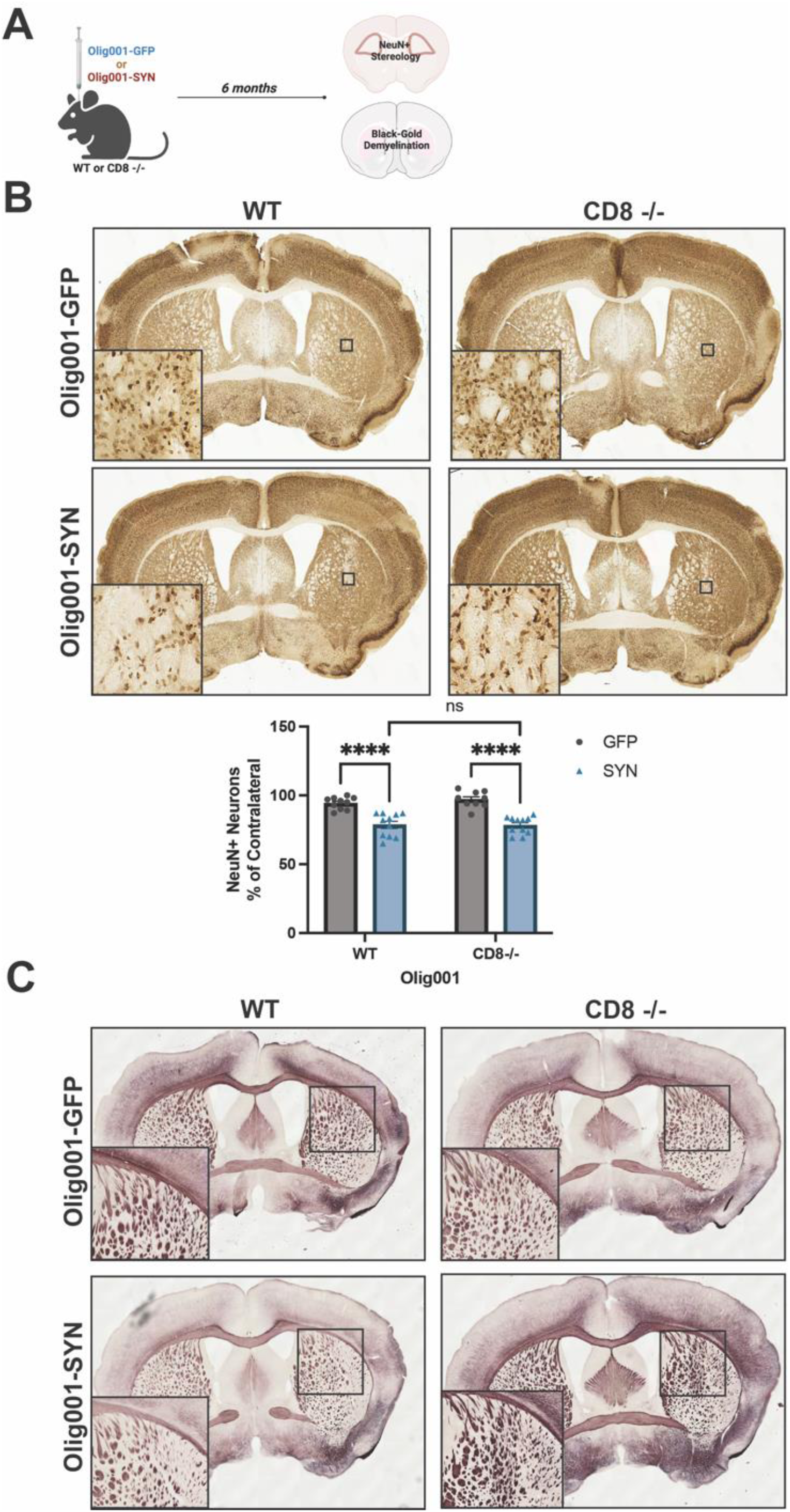
Neurodegeneration and demyelination was unchanged in WT and CD8 knockout mice six months following Olig001-SYN transduction. (**A**) WT and CD8 knockout were injected with either Olig001-GFP or Olig001-SYN, and six months post Olig001-GFP or Olig001-SYN injection tissue was collected and assessed for neurodegeneration and demyelination. (**B**) Representative brightfield images of NeuN+ neurons in the dorsolateral striatum. Small black boxes represent the zoomed in areas (bottom left corner of their corresponding image). Below is the quantification of unbiased NeuN+ neurons within the dorsolateral striatum of WT or CD8 −/− mice injected with either Olig001-GFP or Olig001-SYN. Mean values +/− SEM are plotted, two-way ANOVA with Tukey post hoc for significance, ns= no significance, and **** p<0.0001. N=10-12 for each group, calculated outliners were excluded from final analysis. (**C**) Representative images of Black-Gold stains of WT and CD8 −/− mice injected with either Olig001-GFP or Olig001-SYN. Zoom inserts of affected areas are located to the bottom left of their representative image. For Black-Gold stains, n=8 for each group, only representative images are shown. Mixed groups of male and female mice were used for all experiments.

**Figure 3:**
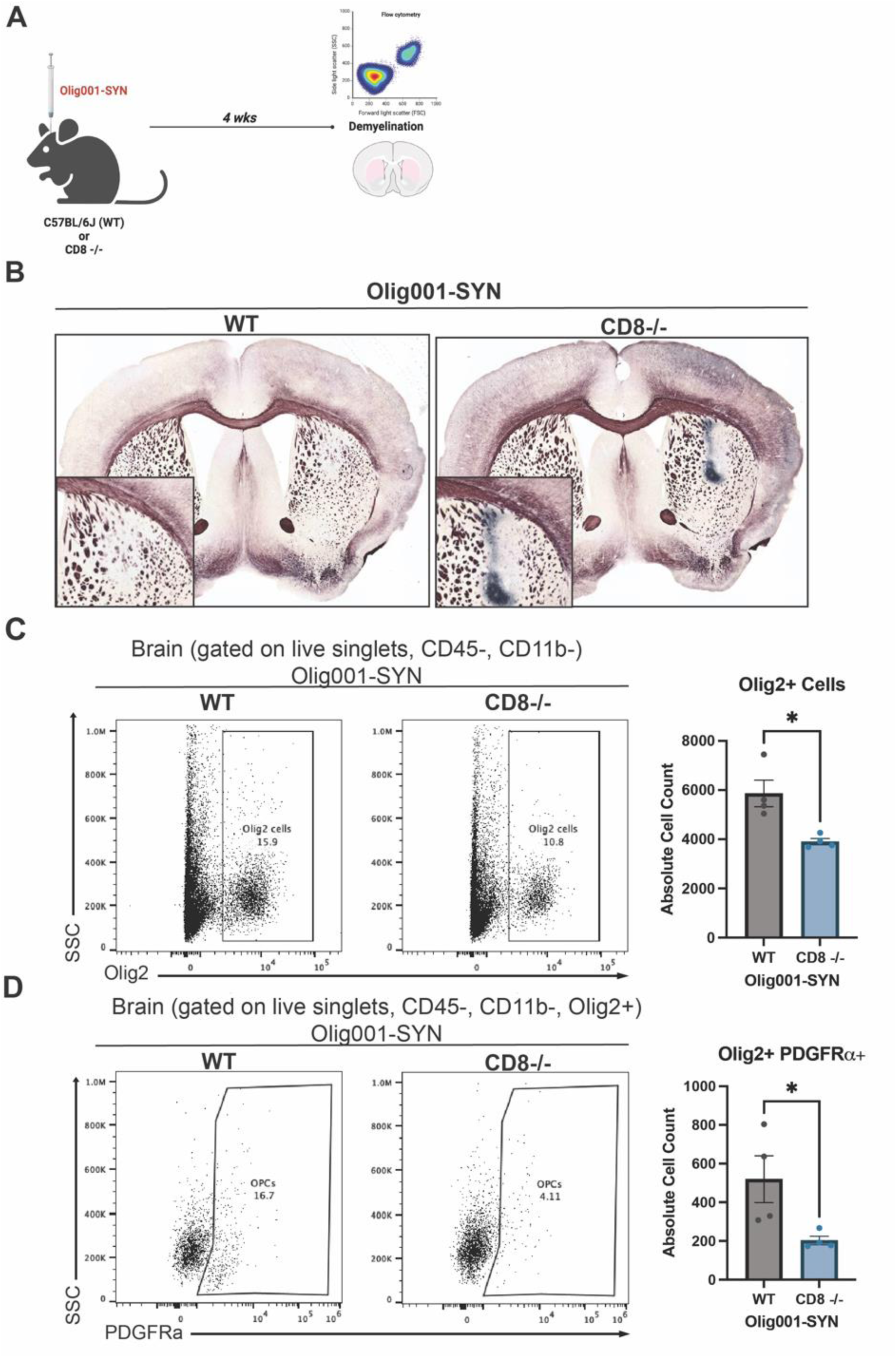
CD8 knockout enhanced demyelination in the Olig001-SYN mouse model. (**A**) WT and CD8 knockout mice at 8-12 weeks were injected with Olig001-SYN, and after four weeks tissue was collected. For flow cytometry experiments, n=4 or 5 were used for each group. For Black-Gold stains, n=5 for each group. (**B**) Representative Black-Gold images of WT and CD8 knockout tissue with GCI pathology. Higher magnification inserts with a magnification of 20x of areas of demyelination are located at the bottom left of their corresponding images. (**C**) Flow cytometry dot plots of Olig2+ cells in WT and CD8−/− mice injected with Olig001-SYN. Quantification of flow cytometry is to the right of dot plot. Mean values +/− SEM are plotted, non-parametric Wilcoxon test, *p < 0.05. (**D**) Flow cytometry dot plots and quantification to the right of OPCs (gated on singlets, live cells, CD11b-, CD45-, Olig2+, PDGFRa+). Mean values +/− SEM are plotted, non-parametric Wilcoxon test, *p < 0.05. For flow cytometry experiments, n = 4 per group, two mice were pooled for each sample. Both male and female mice were used for all experiments.

## Data availability

The authors affirm that the findings of this manuscript are supported by the data therein. Additional information can be requested from the corresponding author.

## Results

### α-syn overexpression in oligodendrocytes enhances CD8+ T cells in the brain parenchyma

In human post-mortem MSA tissue, we have previously reported an increase in CD8+ T cells within the putamen and substantia nigra[40]. Based on these studies, we aimed to explore the distribution of CD8+ T cells within the Olig001-SYN mouse model. WT mice aged 8–12 weeks were bilaterally injected with Olig001-GFP or Olig001-SYN into the dorsolateral striatum. After 4 weeks, the striatum and dura mater meninges were collected and assessed for CD8+ T cells via immunohistochemistry and flow cytometry (Figure 1A). CD8+ T cells infiltrated into the dorsolateral striatum of the brain in response to oligodendrocyte-mediated expression of α-syn (Figures 1B and D). Within the meninges, CD8+ T cells were decreased in the dura mater (Figure 1C). Our results indicate that α-syn overexpression in oligodendrocytes results in increased CD8+ T cells in the brain parenchyma and reduced CD8+ T cells in the dura meninges, indicating an active α-syn-driven neuroinflammatory response.

### The genetic knockout of CD8+ T cells does not prevent neurodegeneration and demyelination in the Olig001-SYN mouse model

Our previous studies detected early neurodegeneration in the dorsolateral striatum at six months post Olig001-SYN injection[7]. To determine whether CD8+ T cells influence neurodegeneration, WT and CD8 knockout mice at 8-12 weeks of age were unilaterally injected with either Olig001-GFP or Olig001-SYN in the dorsolateral striatum. After six months, tissue was harvested for unbiased stereology and Black-Gold myelin staining to determine neurodegeneration and demyelination (Figure 2A). All NeuN+ neurons within the sample were counted in an unbiased manner within the dorsolateral striatum. No changes between the ipsilateral and contralateral sides were detected in the Olig001-GFP WT group. Unlike the Olig001-GFP WT group, there was a significant decrease in NeuN+ neurons on the ipsilateral side compared to the contralateral side in Olig001-SYN WT mice (Figure 2B). A similar decrease in NeuN+ neurons was observed in the Olig001-SYN CD8 knockout mice at six months post α-syn transduction (Figure 2B). No differences were detected between Olig001-SYN injected WT and CD8 knockout mice (Figure 2B), suggesting that the absence of CD8+ T cells does not enhance neurodegeneration in the Olig001-SYN mouse model. Additionally, no observational differences in myelin were detected between Olig001-SYN WT and CD8 knockout mice (Figure 2C). Combined, these results indicate that CD8+ T cells do not attenuate neurodegeneration and demyelination within the Olig001-SYN mouse model at a six-month timepoint.

### CD8 knockout enhances demyelination in the Olig001-SYN mouse model at 4 weeks post model transduction

Demyelination is a progressive pathology in MSA that precedes neurodegeneration[19, 21]. In the Olig001-SYN rodent model, significant demyelination is detected at 4 weeks post Olig001-SYN transduction[7, 26, 40]. To determine the role of CD8+ T cells in demyelination, CD8 knockout and WT (control) mice aged 8-12 weeks old were transduced with Olig001-SYN in the dorsal lateral striatum. 4 weeks post transduction, tissue sections were assessed for myelin integrity with a Black-Gold stain, and oligodendroglia were observed with flow cytometry (Figure 3A). Utilizing the Black-Gold stain, significant demyelination was detected in both WT and CD8 knockout mice, notably with larger lesion areas present in the striatum of CD8 knockout mice (Figure 3B). The areas of GCI pathology were unchanged between WT and CD8 knockout mice when α-syn was overexpressed in oligodendrocytes (Supplemental Figure 1). To further investigate oligodendroglia numbers, flow cytometry was used to determine the number of oligodendrocyte progenitor cells (OPCs; Olig2+ PDGFRa+) located in the striatum of both WT and CD8 knockout mice. Genetic knockout of CD8+ T cells led to a decrease in total Olig2+ cells (Figure 3C), suggesting a loss of oligodendroglia. Furthermore, there was a significant decrease in the number of OPCs (Figure 3D). In mice lacking CD8+ T cells, we observed a significant loss of Olig2+ cells and OPCs, indicating a loss of oligodendroglia. Together, these results demonstrate that the lack of CD8+ T cells during MSA pathogenesis enhances demyelination and the loss of oligodendrocytes.

### Genetic knockout of CD8+ T cells results in enhanced neuroinflammation in the Olig001-SYN mouse model

Demyelination is accompanied by myeloid activation and infiltrating T cells during MSA[40]. Previous studies have suggested that in neurodegenerative diseases, CD8+ T cells can modulate the neuroinflammatory response by secreting cytokines that amplify inflammation and induce apoptosis[11, 14, 15]. Based on this evidence, we sought to investigate the role of CD8+ T cells in α-syn-induced neuroinflammation. Both male and female WT and CD8 knockout mice were transduced with Olig001-SYN at 8-12 weeks of age. Four weeks post transduction, neuroinflammation within the striatum was assessed with flow cytometry and IHC (Figure 4A). With flow cytometry, increased myeloid (CD45+, CD11b+) and lymphocyte (CD45+, CD11b-) populations were detected in the CD8 knockout mice at four weeks post Olig001-SYN transduction (Figure 4B). Within the myeloid population, there were no changes in the number of microglia in the dorsolateral striatum, however there was in increase in infiltrating Ly6C^hi^ monocytes into the parenchyma of CD8 knockout mice (Figure 4C). Although the amount of microglia did not change between groups, there was increased MHCII+ expression on microglia in Olig001-SYN CD8 knockout mice (Figure 4D and E). Taken together, these results suggest an enhanced myeloid neuroinflammatory response in CD8 knockout mice, highlighting a potential role for CD8+ T cells in modulating neuroinflammation.

**Figure 4:**
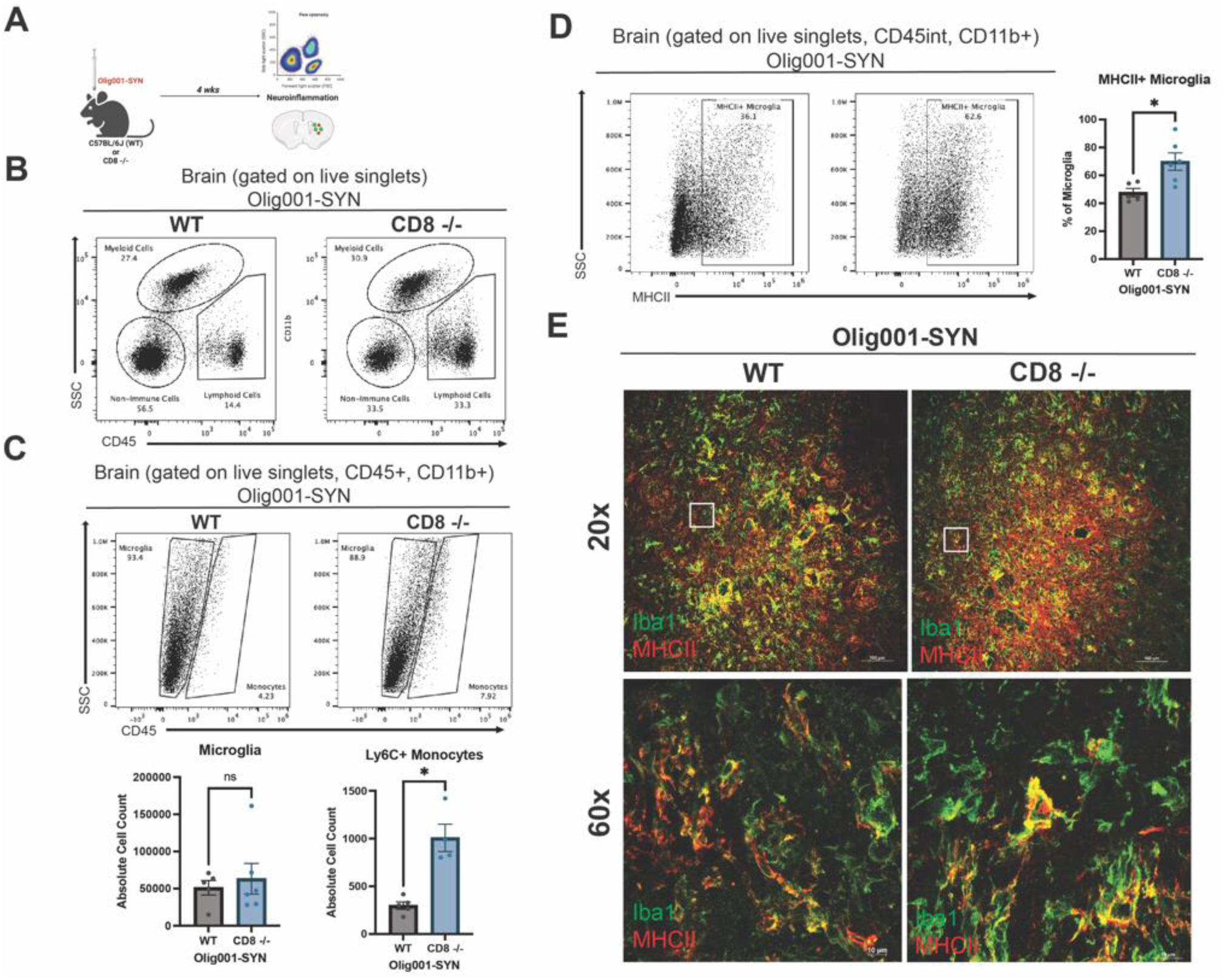
CD8 knockout enhances MHCII expression and infiltration of Ly6Chi monocytes in mice transduced with Olig001-SYN. (**A**) WT and CD8 knockout mice at 8 weeks old were injected with Olig001-SYN. At four weeks post injection, tissue was collected for flow cytometry and immunohistochemistry. For flow cytometry experiments, n=4 or 5 were used per group (2 mice were pooled for each sample). For immunohistochemistry, n=4 per group. (**B**) Dot plots of the general gating strategy to assess for myeloid cells and lymphocytes. (**C**) Representative flow cytometry dot plots of microglia (gated on singlets, live cells, CD11b+, CD45^int^), monocytes (gated on singlets, live cells, CD11b+, CD45^hi^), and Ly6C+ monocytes (gated on singlets, live cells, CD11b+, CD45+, Ly6C+) in the striatum of WT and CD8 knockout mice with transduced with Olig001-SYN; quantification of microglia and Ly6C+ monocytes are below the dot plots. Mean values +/− SEM are plotted, non-parametric Wilcoxon test, ns= no significance, *p < 0.05. (**D**) Flow cytometry histograms of MHCII+ microglia (gated on singlets, live cells, CD11b+, CD45^mid^, MHCII+) in the striatum of WT and CD8 knockout mice injected with Olig001-SYN; the quantification of MHCII+ microglia are to the right. Mean values +/− SEM are plotted, non-parametric Wilcoxon test, ns= no significance, *p < 0.05. (**E**) Representative images show Iba1+ cells (green) in both WT and CD8 knockout mice, and enhanced MHCII+ (red) in Olig001-SYN transduced CD8 knockout mice. White boxes indicate areas of higher magnification below. Low magnification confocal images (scale bar = 100 μm; top panel images), High magnification confocal images (bottom panel; scale bar = 10 μm). Both male and female mice were used in this study.

Lymphocyte populations (CD45+, CD11b-) were also elevated in the striatum of CD8 knockout mice transduced with Olig001-SYN. At four weeks post Olig001-SYN injection, CD8 knockout had increased infiltration of CD4+ T cells into the brain parenchyma (Figure 5B and C). Additionally, CD4+ T cells were located around areas of pSer129+ GCI pathology (Figure 5C). pSer129+ GCI pathology was unchanged in WT and CD8 knockout mice at four weeks (Supplemental Figure 1). No changes were observed in the general B cell population (Figure 5D) and IgG deposition (Figure 5E). Although there were no changes in infiltrating B cells between WT and CD8 knockout mice, we have previously reported that CD4+ T cells are critical in driving neuroinflammation in the Olig001-SYN mouse model. The infiltration of CD4+ T cells around areas of GCI pathology in the CD8 knockout mice imply an amplified neuroinflammation response.

**Figure 5:**
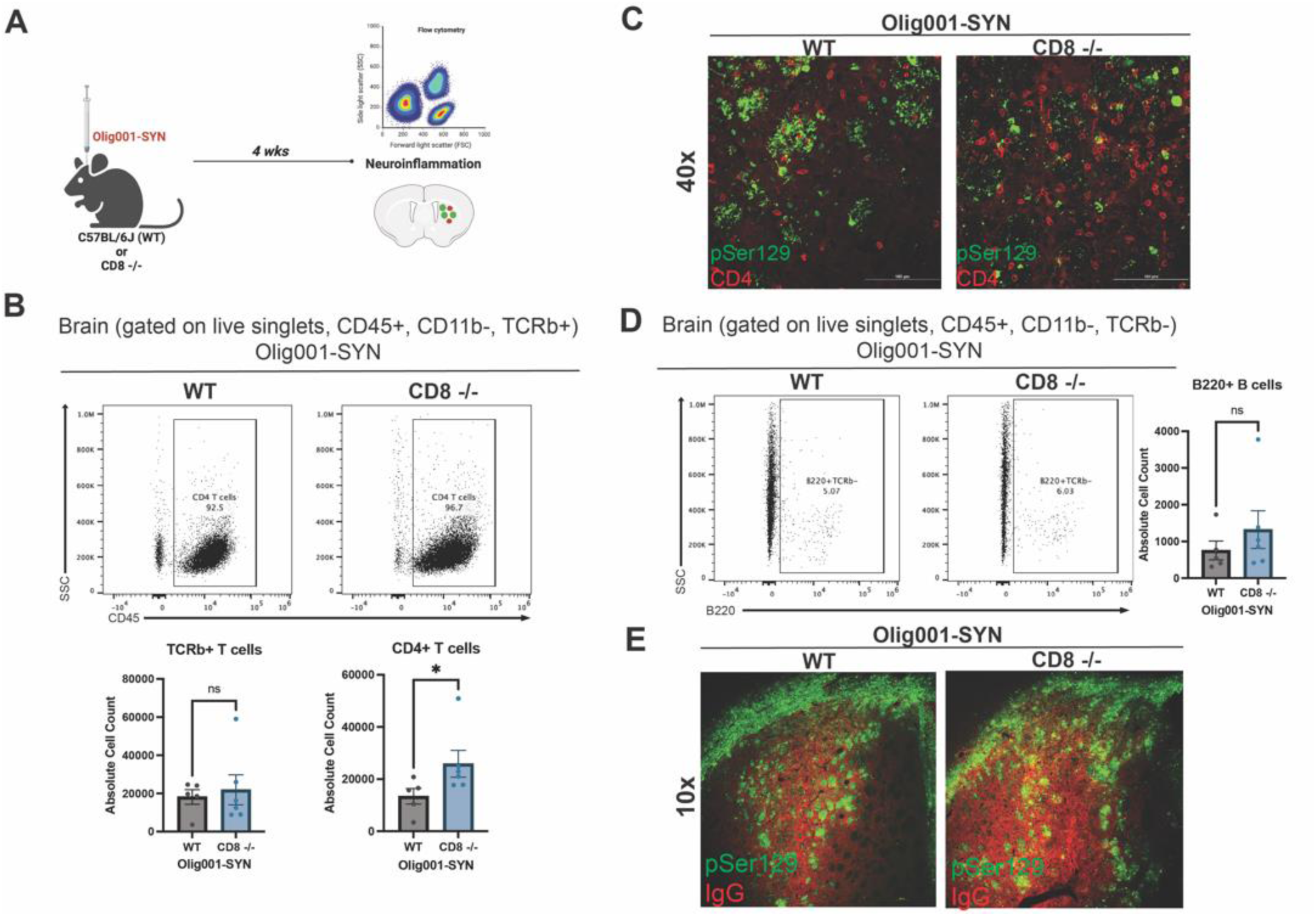
CD8 knockout mice show enhanced infiltration of CD4+ T cells and unchanged B cell infiltration and IgG deposition at four weeks post Olig001-SYN transduction. (**A**) Both WT and CD8 knockout mice were injected with Olig001-SYN into the dorsolateral striatum, and tissue was collects four weeks later to assess for neuroinflammation with flow cytometry and immunohistochemistry. (**B**) Flow cytometry dot plots of CD4+ T cells (gated on singlets, live cells, CD11b-, CD45+, TCRb+ CD4+) in striatum of WT and CD8 knockout mice with α-syn induced neuroinflammation; the quantification is to the right of the dot plots. Mean values +/− SEM are plotted, non-parametric Wilcoxn test, ns= no significance, *p < 0.05. n= 4/5 per group, with two mice pooled per sample. (**C**) Representative immunohistochemistry images of CD4+ T cells (red) surrounding α-syn (pSer129; green) in WT and CD8 knockout mice with GCI pathology (scale bar = 100 μm, n=3 per group). (**D**) Flow cytometry dot plots of B cells (gated on singlets, live cells, CD11b-, CD45+, TCRb-, B220+) of WT and CD8 knockout injected with Olig001-SYN, the quantification of the dot plots for B cells is to the right of the dot plots. Mean values +/− SEM are plotted, non-parametric Wilcoxn test, ns= no significance. n= 4/5 per group, with two mice pooled per sample. (**E**) Representative immunohistochemistry images taken at 10x of IgG deposition (IgG; red) in WT and CD8 knockout mice when α-syn (pSer129; green) is present. Scale bar = 100 μm. n=3 per group. Both female and male mice were used for all experiments.

### Tissue resident memory CD8+ T cells are present in the Olig001-SYN mouse model

In post-mortem MSA tissue, CD8+ T cells are present; however, their differentiation states are unknown[40]. In another synucleinopathy, PD, CD8+ T cell present in the substantia nigra display markers for tissue residency and memory[6, 11]. Additionally, a similar tissue resident memory phenotype is present on CD8+ T cells within white matter areas of MS tissue[9]. To further understand the type of CD8+ T cells present during disease, we sought to determine the state CD8+ T cells were in within the dorsolateral striatum. To understand the phenotype of CD8+ T cells in the Olig001-SYN mouse model, WT mice were bilaterally injected with 2uL of either Olig001-GFP or Olig001-SYN into the striatum. Pathology developed for four weeks, and the striatum was collected for spectral flow cytometry (Figure 6A). There were a total of 2921 CD8+ T cells, and 7588 CD8+ T cells present in the Olig001-GFP and Olig001-SYN groups respectively (Figure 6B). Utilizing spectral flow cytometry, the co-expression of CD8+ with various markers were identified (Figure 6C). CD8+ T cells present within the striatum of Olig001-SYN transduced mice expressed markers associated with tissue residency (CD103+), tissue retention (CD69+), early memory (CD44+ and CD27+), exhaustion (PD-1) (Figure 6H), tissue honing (CXCR3 and CCR5), tissue-resident memory (CXCR6+), and senescence (KLRG1) (Figure 6C). Again, using unbiased analysis, we observed that the PD-1+, CD44+, CXCR6+, and CD103+ CD8+ T cell populations had a higher expression in the Olig001-SYN group when compared to Olig001-GFP controls (Figure 6D). Upon further investigation, 9 unique populations of CD8+ T cells were identified between Olig001-GFP and SYN (Figure 6E). Although there was an overall increase in CD8+ T cells in the Olig001-SYN group, there were larger frequencies of CD8+ T cells in clusters 1 (CD103+, CD69+, KLRG1+), 3 (CD103+, PD-1+, CD44+, CCR5+), and 5 (CD103+, CD44+, KLRG1+, PD-1+, CD27+, CD69+, CXCR3+, CXCR6+) coupled with decreased frequencies in clusters 2 (KLRG1+, CD27+, CD69+, CXCR3+), 6 (CD62L+), and 8 (CD27+, CD69+, CXCR3+, CCR5+) (Figure 6F). In the Olig001-SYN-injected mice, there was an increase in CD103+, PD-1+ (Figure 6G-H), CD44+, and CXCR6+ CD8+ T cells (Figure 6G). Notably previous research has shown that CXCR6+ CD8+ T cells that co-express markers for exhausted (PD-1+) tissue-resident (CD103+, CD69+) memory (CD44+, CD27+) are vital in suppressing neuroinflammation and preventing cognitive decline in the 5xFAD mouse model[36]. The results presented here suggest that CD8+ T cells are increased in the Olig001-SYN mouse model and express markers of tissue resident memory (CD103+, CD69+, CD44+, CD27+, CXCR6+) cells.

**Figure 6:**
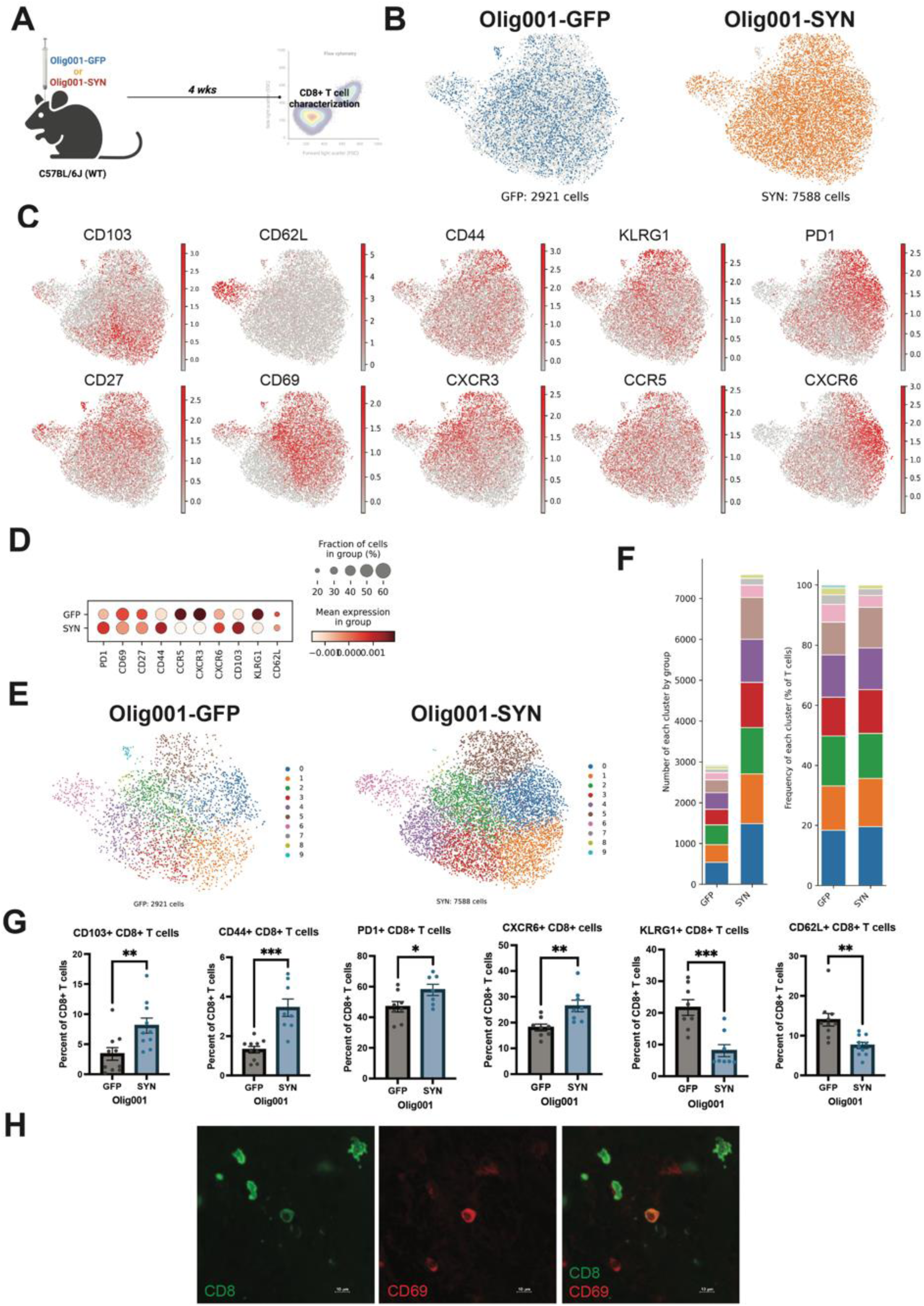
Olig001-SYN transduction enhances CD8+ T cell subsets in the striatum of mice at four weeks. (**A**) WT mice were bilaterally injected at 8-12 weeks old with Olig001-GFP (control) or Olig001-SYN in the dorsal lateral striatum harvested four weeks later for spectral flow cytometry. (**B**) UMAP display the total amount of CD8+ T cells between Olig001-GFP and SYN. (**C**) UMAP of CD8+ T cells and cell surface markers associated with tissue resident memory in Olig001-SYN inject WT and CD8 knockout mice. (**D**) Group comparison of the CD8+ T cells tissue resident memory markers between Olig001-GFP and SYN. (**E**) UMAP of Olig001-GFP and SYN CD8+ T cells identifying 9 clusters associated with unique subpopulations of CD8+ T cells. (**F**) The number of cells within each cluster between Olig001-GFP and SYN (left), and the frequency of each cluster of CD8+ T cells between Olig001-GFP and SYN (right). (**G**) quantification spectral flow cytometry of the following CD8+ T cells (gated on singlet, live cells, CD11b-, CD45+, TCRb+, CD8+): CD103+, CD44+, PD-1+, CXCR6+, KLRG1+, and CD62L+. Mean values +/− SEM are plotted, parametric unpaired t-test, *p < 0.05, **p<0.01, *** p <0.001. All spectral flow cytometry experiments had a n=10/group, each point represents one sample and 2 mice were pooled per sample. Both male and female mice were used. All UMAPs presented were generated using metadata from the spectral flow cytometry experiment. (**H**) Representative images of CD8+ (green) and CD69+ (red) T cells in Olig001-SYN injected mice. Scale bar is set at 10 μM. Both male and female mice were used for all experiments.

## Discussion

In this study, we identified tissue-resident memory CD8+ T cells in the parenchyma of the dorsolateral striatum when α-syn was overexpressed in oligodendrocytes, modeling human MSA pathology in mice (Figure 1). To determine the role of CD8+ T cells during MSA pathogenesis, we utilized CD8+ T cell knockout mice. Six months after Olig001-SYN transduction, demyelination and neurodegeneration were unchanged in CD8 knockout mice indicating CD8 T cells do not attenuate α-syn-mediated MSA pathology (Figure 2). However, when evaluating consequences of Olig001-SYN transduction at an intermediate four-week timepoint we found that CD8 knockout mice displayed enhanced demyelination and neuroinflammation (Figures 3, 4, and 5). Utilizing unbiased spectral flow cytometry, we identified that the CD8+ T cells present were differentiated into tissue-resident memory T cells (Trm) (Figure 6). This study demonstrates that CD8+ T cells are not critical in driving disease; however, without CD8+ T cells, early enhanced pathology will occur. Additionally, this study is the first to suggest the presence of CD8+ Trm cells in MSA.

Previous studies have reported that CD8+ Trm cells are located in white matter lesions in demyelinating diseases like MS[9]. In human MSA post-mortem tissue, we have previously reported an increase in CD8+ T cells within the putamen and substantia nigra[40]. Literature has also shown that CD8+ T cells are retained in dura and leptomeningeal spaces, and upon a stimulus in the brain, will infiltrate into the parenchyma and execute effector functions[1]. Our results potentially suggest a similar trafficking of CD8+ T cells into the brain that has been described in other studies as α-syn overexpression results in enhanced parenchymal CD8+ T cells but decreased dural CD8+ T cells (Figure 1B and C). Previous studies investigating CD8+ T cells in subcortical white matter areas have identified large population of CD8+ Trm cells within the corpus callosum of healthy individuals [35]. In PD, early-stage post-mortem tissue has shown that the infiltration of CD8+ Trm cells occurs when α-syn pathology was only in the olfactory bulb and not the substantia nigra[11]. Interestingly, the CD8+ T cells identified in early-stage PD study also shared markers for tissue residency (CD103 and CD69) indicating the presence of CD8+ Trm cells may be age and disease dependent. Similarly, within the striatum, we found CD8+ T cells also expressed markers of tissue residency (Figure 6).

Neurodegeneration was assessed six months after model induction in WT and CD8 knockout mice, using unbiased stereology to analyze NeuN+ neurons within the dorsolateral striatum (Figure 2A). Our results indicated that neurodegeneration was not attenuated in the Olig001-SYN CD8 knockout mice when compared to their Olig001-GFP control mice (Figure 2B). The percentage of NeuN+ cell loss was not statistically significant between WT and CD8 knockout mice transduced with Olig001-SYN (Figure 2B), despite similar levels of α-syn expression (Supplemental Figure 1). The Black-Gold stain show that demyelination was unchanged between WT and CD8 knockout mice with α-syn overexpression in oligodendrocytes (Figure 2C). Although our results indicate that neurodegeneration and demyelination are not mediated by CD8+ T cells, previous studies have shown that the genetic knockout of Tbet-dependent IFNγ does lead to attenuated neurodegeneration at six months post oligodendrocyte mediated α-syn induction[7]. CD8+ T cells can produce IFNγ [14], however, our group has demonstrated the main producers of IFNγ during MSA are CD4+ T cells [7]. To determine the effects of the adaptive immune system in driving neurodegeneration during MSA, more long studies focused on CD4+ T cells are warranted.

Early infiltration of CD8+ T cells into the brain parenchyma can occur in MS, PD, and with age [9, 11, 35]. Demyelination is a critical pathological hallmark that precedes neurodegeneration during MSA [19, 21]. To determine if CD8+ knockout protects against demyelination during MSA at an early-stage time point, we conducted flow cytometry and immunohistochemistry four weeks after Olig001-SYN induction in WT and CD8 knockout mice and assessed demyelination (Figure 3). We found that demyelination was not attenuated but enhanced in the CD8 knockout mice transduced with Olig001-SYN compared to Olig001-SYN injected WT mice (Figure 3B). Flow cytometry revealed that Olig2+ cells (a marker for oligodendroglia) and PDGFRa (a marker for oligodendrocyte progenitor cells) were decreased in Olig001-SYN CD8 knockout mice when compared to WT mice transduced with Olig001-SYN (Figure 3B and C). Characterization of the oligodendrocytes within the Olig001 mouse model displayed no changes in general oligodendroglia and their progenitor cells (OPCs) between the GFP and SYN groups (Supplemental Figure 2). The demyelination data presented in Figure 3 (a four-week timepoint) and Figure 2C (a six-month timepoint) suggest that the lack of CD8+ T cells will cause demyelination early in disease pathogenesis. There is growing evidence in the PD field to suggest that CD8+ T cells infiltrate early into the brain when α-syn is present[11]. Additionally, in MS, another demyelination disease, CD8+ Trm cells are present in white matter lesion in early disease progression[9]. The direct interaction between CD8+ T cells and oligodendrocytes is not known in MSA, but previous studies have shown that IFNγ signaling is upstream of demyelination and ultimately neurodegeneration[7]. The direct interactions of oligodendrocyte dysfunction with the neuroinflammatory response during MSA are largely unknown[18, 19, 21, 25]. Overall, the data presented in Figure 2 suggest that CD8+ T cells do enhance demyelination at an early timepoint however the mechanism in which the lack of CD8+ T cells contributes to demyelination is unknown. Due to neuroinflammation and demyelination occurring before neurodegeneration in MSA, future studies are needed to understand if CD8+ T cells are vital for oligodendrocyte health in earlier time points.

In other neurodegenerative disease like AD and PD, CD8+ T cells contribute to the neuroinflammatory response by secreting proinflammatory cytokines that can induce apoptosis[5, 6, 12, 39]. To determine if CD8+ T cells were proinflammatory around areas of demyelination, flow cytometry and immunohistochemistry (Figure 4A and 5A) was used at the 4-wek timepoint in Olig001-SYN injected CD8 knockout mice. Indeed, we found increased MHCII expression (Figure 4D and E) and infiltration of monocytes (Figure 4C), indicating an enhanced myeloid response. Additionally, in α-syn treated CD8 knockout mice, CD4+ T cells were increased in the parenchyma (Figure 5B and C), with little change observed in infiltrating B cells (Figure 5D) and IgG deposition (Figure 5E). Our previous studies, along with others, have shown myeloid activation and infiltration of T cells are critical in driving neuroinflammation in various mouse models of AD and PD[20, 40, 41]. Recent literature has highlighted the role of CD8+ T cells in regulating the proinflammatory functions of CD4+ T cells. In a chronic inflammation model of long COVID, suppressive Ly49+ CD8+ T cells regulate autoreactive CD4+ T cells via IL-10 cytokine secretion [23, 32]. Conversely in a chronic infection and EAE mouse models, CD4+ T cells help maintain CD8+ Trm cells within the brain[15, 29, 32, 38]. Our group has showed that CD4+ T cells and their production of IFNγ are critical in mediating neurodegeneration and demyelination in the Olig001-SYN mouse model[7, 40]. Although α-syn-induced neuroinflammation and demyelination were attenuated when CD4+ T cells were not present[40], CD8+ T cells were not investigated. Due to the importance of CD4+ T cells in driving MSA and the interplay that occurs between CD4+ and CD8+ T cells, further studies are needed to determine the interaction of these two important cell types in driving neuroinflammation.

In the Olig001-SYN mouse model of MSA, CD8+ T cells contribute to the demyelination and neuroinflammation (Figure 3, 4, and 5) at four weeks, however the specific phenotype of these CD8+ T cells during MSA are unknown. Utilizing spectral flow cytometry, we characterized CD8+ T cells in the dorsolateral striatum in Olig001-SYN transduced mice. Four weeks post transduction, we found that CD8+ T cells are increased in the brain parenchyma surrounding areas of GCI pathology (Figure 1) and are co-expressed with various markers for tissue adhesion (CD103), tissue retention (CD69), chemokine receptors associated with tissue homing (CCR5, CXCR6, CXCR3), and early memory (CD44, CD27). The unbiased group comparison between Olig001-GFP and Olig001-SYN displayed increased expression of PD-1, CD44, CXCR6, and CD103, which were verified with FlowJo analysis (Figure 6D and H). Interestingly, our spectral flow cytometry results also highlighted a decrease in KLRG1 (Figure 6D and G), a marker associated with T cell senescence and terminal differentiation[16, 17]. Additionally, the decrease in KLRG1 is also coupled with the increase of PD-1, a T cell exhaustion marker[2, 14]. We also identified an increased CXCR6+ CD8+ T cell population within the Olig001-SYN mouse model (Figure 6C and D). Recent findings have also identified exhausted (PD-1+) CXCR6+ CD8+ T cells as protective in the 5xFAD and the APP mouse models of AD [36]. The exact mechanism of how CXCR6+ CD8+ T cells are protective within the 5xFAD and APP is unknown, however there is evidence to suggest CXCR6+ CD8+ T cells in the 5xFAD model will interact with disease-associated microglia [36]. The findings from our study strongly suggests the presence of CD8+ Trm cells in the brain parenchyma during MSA, however future studies are warranted to understand the specific role CD8+ Trm cells during MSA.

The results provided by the current study help to highlight the complexity of CD8+ T cells not only in MSA, but in neurodegenerative diseases in general. T cells, including CD8+ T cells, become activated upon encountering antigens through MHCI expression[14, 15]. After clonal expansion, T cells transition into memory T cells, which are long-lived primed cells capable of residing throughout the body, including the brain[14, 15]. In neurodegenerative diseases like AD, PD, and MS, clonally expanded CD8+ T cells expressing markers of resident memory have been detected in CSF[2, 9, 12, 14, 39]. Additionally, CD103+ CD69+ CD8+ T cells, markers of tissue retention and residence, have been detected in the CSF of AD, PD, and MS patients[9, 10, 12, 39]. In a recent PD study, evidence of CD8+ Trm cells within the parenchyma in the early stages of PD pathogenesis have been found[11]. CD8+ Trm cells have also been detected in the white matter lesions in post-mortem MS tissue[9]. Our previous work showed an increase of CD8+ T cells within MSA post-mortem tissue[40]; however, it is unknown if they are clonally expanded and if they express markers of tissue resident memory. The results yielded from this study identified CD103+ CD69+ CD44+ CD8+ T cells within the striatum, suggesting there are early CD8+ Trm cells in the parenchyma in MSA. Although clonal expansion is vital for the formation of Trm cells, this process has not yet been demonstrated in this model. Future studies are necessary in human post-mortem tissue to determine whether CD8+ T cells are clonally expanded in the putamen of MSA patients and if they exhibit markers for tissue residency and memory.

Altogether, our findings show that CD8+ T cells are not critical for mediating MSA pathology in Olig001-SYN mice, but may be important in modulating early disease pathology. Genetic knockout of CD8+ T cells resulted in enhanced demyelination and a reduction in Olig2+ cells and OPCs within the lesion area. Additionally, knockout of CD8+ T cells enhanced CNS microglia activation and antigen presentation, the infiltration of proinflammatory monocytes and CD4+ T cells. Through spectral flow cytometry and unbiased analysis, we observed an increased expression of markers associated with tissue residency, memory, and exhaustion in CD8+ T cells within the Olig001-SYN model. Taken together, our results highlight a potential role of CD8+ Trm cells in modulating neuroinflammation and demyelination during MSA disease progression.

## Abbreviations

α-syn: alpha-synuclein
CNS: Central Nervous System
Trm: tissue resident memory T cell
APC: antigen presenting cells
GCI: glial cytoplasmic inclusions
IFNγ: Interferon gamma
MHCI: major histocompatibility complex I
MHCII: major histocompatibility complex II
MSA: multiple system atrophy
AD: Alzheimer’s disease
PD: Parkinson’s disease
OPC: oligodendrocyte progenitor cell

## Declarations

## Acknowledgements

The work in these studies was generously supported by NIH/NINDS R01NS107316 (Harms, PI). We are also grateful for the experimental input, discussion, and key reagents provided by Chander Raman. We are also thankful for Vidya Sagar Hanumanthu and the UAB Comprehensive Flow Cytometry Core for their assistance in flow cytometry experiments, and Anna Stoll for critical feedback on the manuscript. Experimental design figures were generated with BioRender.

## Author’s Contributions

NJCS and ASH designed the study and wrote the manuscript. NJCS was responsible for stereotaxic surgeries. NJCS, GMC, JMW, AZ, and YTY contributed towards tissue collection and executing flow cytometry experiments. MAA developed the spectral flow cytometry analysis pipeline. MAA and DJT applied the spectral flow cytometry pipeline to the Olig001-GFP and SYN data. NJCS conducted all IHC stains. NJCS assessed NeuN+ stereology. NJCS and ASH analyzed flow cytometry, IHC, and stereology data. NJCS and ASH drafted and constructed the manuscript and figures. FPM developed and optimized the Olig001 vector model. FPM, JK, DJT provided feedback and edits to the manuscript. All authors read and approved the final manuscript.

## The Supplementary material

