## Supplemental Figures for "Tissue resident memory CD8+ T cells are present but not critical for demyelination and neurodegeneration in a mouse model of multiple system atrophy"

**Supplemental Figures**
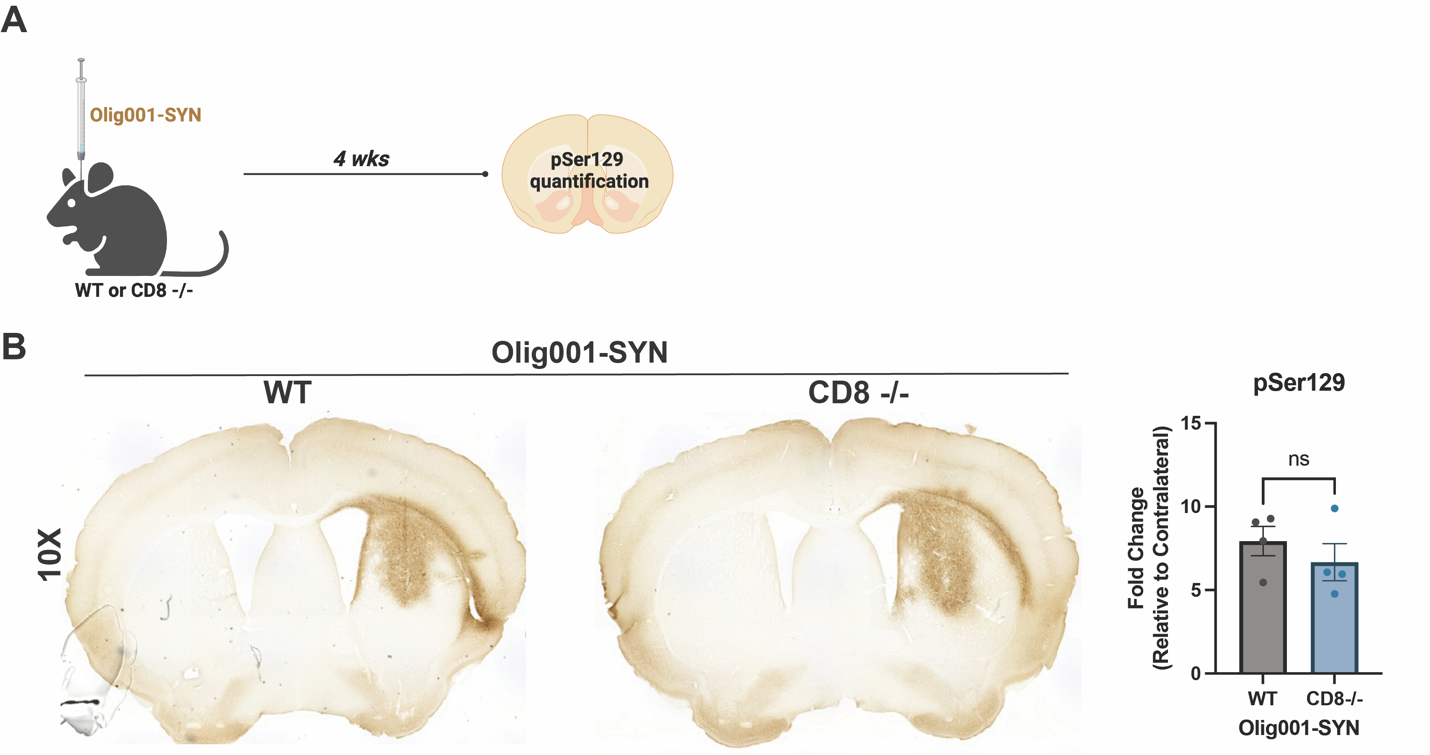

**Supplemental Figure 1: pSer129+ GCI pathology was unchanged in WT and CD8 knockout mice 4 weeks post Olig001-SYN transduction.** (**A**) WT and CD8 knockout were unilaterally injected with Olig001-SYN, and at 4 weeks tissue was collected to assess for pSer129 pathology. (**B**) Representative images of DAB pSer129 stains of WT and CD8 knockout mice injected with Olig001-SYN. Mean values +/- SEM are plotted, non-parametric Wilcoxon test, ns=not significant. n=4 for each group, only representative images are showed. Mix group of both male and female mice were used in each group.

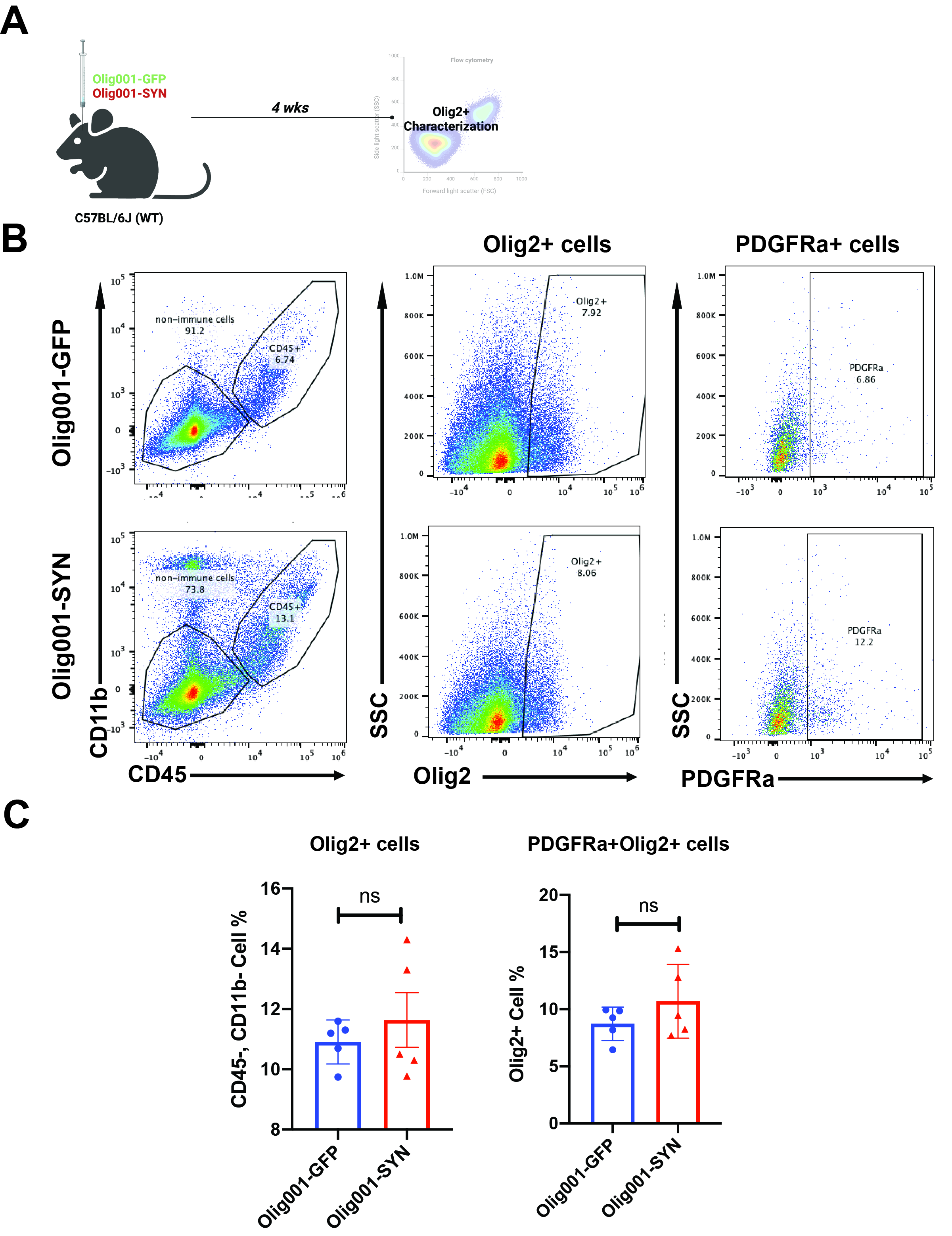
 **Supplemental Figure 2: Oligodendroglia characterization in the Olig001-SYN mouse model.** (**A**) Both female and male WT mice were injected with either Olig001-GFP or Olig001-SYN at 8-12 weeks of age. After 4 weeks, tissue was collected for flow cytometry to assess for oligodendroglia. (**B**) Flow cytometry plots of non-immune cells (gated on singlets, live cells, CD45-, CD11b-), Olig2+ cells in the middle panel (gated on singlets, live cells, CD45-, CD11b-, Olig2+), and OPCs in the right panel (gated on singlets, live cells, CD45-, CD11b-, Olig2+, PDGFRa+). (**C**) Quantification of flow cytometry plots of Olig2+ and Olig2+ PDGFRa+ cells. Mean values +/- SEM are plotted, non-parametric Wilcoxon test, ns=not significant. For flow cytometry experiments, n = 5 per group, two mice were pooled for each sample. Both male and female mice were used for all experiments.
